# ATP-dependent force generation and membrane scission by ESCRT-III and Vps4

**DOI:** 10.1101/262170

**Authors:** Johannes Schöneberg, Shannon Yan, Maurizio Righini, Mark Remec Pavlin, Il-Hyung Lee, Lars-Anders Carlson, Amir Houshang Bahrami, Daniel H. Goldman, Xuefeng Ren, Gerhard Hummer, Carlos Bustamante, James H. Hurley

## Abstract

The ESCRTs catalyze reverse-topology scission from the inner face of membrane necks in HIV budding, multivesicular endosome biogenesis, cytokinesis, and other pathways. We encapsulated a minimal ESCRT module consisting of ESCRT-III subunits Snf7, Vps24, and Vps2, and the AAA^+^ ATPase Vps4 such that membrane nanotubes reflecting the correct topology of scission could be pulled from giant vesicles. Upon ATP release by photo-uncaging, this system was capable of generating forces within the nanotubes in a manner dependent upon Vps4 catalytic activity, Vps4 coupling to the ESCRT-III proteins, and membrane insertion by Snf7. At physiological concentrations, single scission events were observed that correlated with forces of ~6 pN, verifying predictions that ESCRTs are capable of exerting forces on membranes. Imaging of scission with subsecond resolution revealed Snf7 puncta at the sites of membrane cutting, directly verifying longstanding predictions for the ESCRT scission mechanism.

**One Sentence Summary:** ESCRT-III and Vps4 were reconstituted from within the interior of nanotubes pulled from giant vesicles, revealing that this machinery couples ATP-dependent force production for membrane scission.

## Main Text

Cellular membranes are constantly remodeled in the course of vesicular trafficking, cell division, the egress of HIV, and a host of other processes. Membranes can bud and be severed either towards or away from the cytosol. The latter is referred to as “reverse topology” scission and is catalyzed by the ESCRT machinery, a set of ~18 proteins in yeast and ~28 in mammals (*1*–*4*). The core machinery of membrane scission by the ESCRTs consists of the ESCRT-III protein family. Among those, the most important components for membrane scission are Snf7, Vps24, and Vps2 (*5*, *6*). When recruited to membranes, ESCRT-III proteins assemble into flat spiral discs (*7*–*9*), helical tubes (*7*, *10*, *11*), or conical funnels (*11*–*13*). ESCRT filaments have a preferred curvature (*8*, *9*, *14*). When they are bent to curvatures of higher or lower values, ESCRT filaments act as springs that restore their own shape to the preferred value (*9*, *15*, *16*). This spring-like behavior has led to the prediction that ESCRTs could exert measurable forces upon membranes, which we set out to test.

The AAA^+^ ATPase Vps4 (*17*) is intimately associated with the ESCRT machinery and essential for the membrane scission cycle. Vps4 is recruited to scission sites by Vps2 (*18*, *19*). Vps2 is thought to have a capping role whereby it inhibits Snf7 polymerization (*6*). By recycling Vps2 (*20*), Vps4 promotes Snf7 polymerization. Thus, Vps4 is critical for the recycling of ESCRT-III and the replenishment of the soluble cytoplasmic pool. Early attempts at in vitro reconstitution of ESCRT-mediated budding and scission using giant unilamellar vesicles (GUVs) suggested that the process was independent of Vps4 and ATP (*21*, *22*), except for the final post-scission recycling step. Cell imaging studies (*23*–*28*), however, showed that Vps4 localization peaked prior to scission in HIV-1 budding and cytokinesis, consistent with its direct role in scission upon ATP hydrolysis. A second goal of this study was to determine if Vps4 and ATP hydrolysis were directly involved in membrane scission, as opposed to mere recycling.

We encapsulated the minimal ESCRT-III-Vps4 module containing yeast Snf7, Vps24, Vps2, and Vps4 (referred to hereafter as the “module;” Fig. S1), in POPC:POPS:Biotinyl-PE (80:20:0.1) GUVs at near-physiological ionic strength (~150 mM NaCl; Fig. 1A–D). We used optical tweezers to pull nanotubes extending between the surface of a GUV held by suction on an aspiration pipette and the surface of a streptavidin-coated polystyrene bead held by an optical trap (Fig. 1E, F). To fuel the AAA+ ATPase Vps4, we also encapsulated the caged ATP analog NPE-ATP. An optical fiber was used to UV-illuminate one GUV at a time (Fig. S2) so that experiments could be carried out sequentially on individual GUVs in the same micro-fluidic observation chamber. In control experiments, where all protein components were included except for ATP, UV illumination led to no change in the force exerted on the bead (Fig. 1F, 2B). In similar control experiments omitting only Vps4, UV illumination results in a slight drop in the pulling force (Fig. 1G, 2C), attributed to the generation of two product molecules upon NPE-ATP uncaging. Thus, in the absence of ESCRT activity the membrane nanotube is stable.

**Fig. 1.**
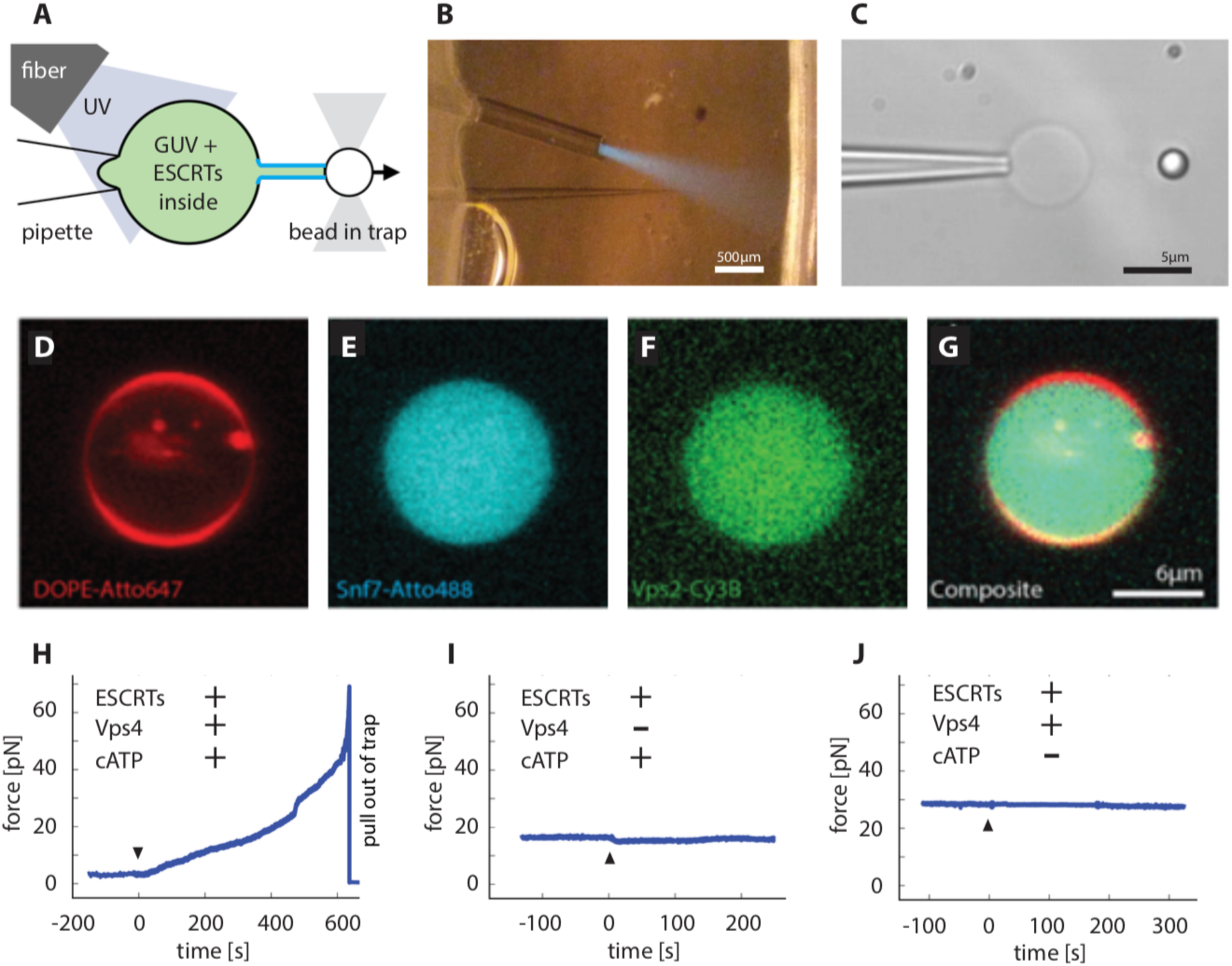
ESCRT-III exerts an ATP dependent force on membrane tubes. A. Schematic drawing: a membrane tube (middle) is pulled out of a micropipette aspirated GUV with a functionalized bead held in an optical trap (see main text), creating reverse curvature topology. Components of the ESCRT machinery and caged ATP are encapsulated in the lumen of GUV. An optical fiber delivers UV light to uncage the ATP inside the vesicle and start the reaction. B. Actual picture of the aspiration pipette (needle-like shape), optical fiber, and UV light cone (blue) carefully configured inside the microfluidic chamber for our experiments (see main text). C. Bright field image of a GUV (center) aspirated by the micropipette (left) and a tube-pulling bead (right). Labeling different components of the ESCRT module (Snf7 in (E), Vps2 in (F), and merged in (G)) revealed a uniform distribution of proteins in the lumen of GUVs (membrane in (D)). H, I, J. Force profile over time detected by optical tweezers on a membrane tube pulled from a GUV that encapsulates components of the ESCRT module. H. ATP uncaging (black arrow head) in the presence of a full ESCRT module leads to a rise in force exerted on the tube, which is connected to the bead held by the optical trap. The large force rise can overcome the trapping strength and eventually pulls the bead out of the trap. I. Control experiment on a full ESCRT module but omitting Vps4. Apart from a minute dip in the force profile, which is due to small osmolarity change upon ATP uncaging, no effects were detected. J. Control experiment on a full ESCRT module but omitting ATP. No change in force measurable.

Initial experiments with ESCRTs were carried out at a super-physiological protein concentration of 2 µM in order to measure effects under overdriven conditions. When ATP was uncaged in the presence of the complete ESCRT module, a large rise in retraction force was indeed observed (Fig. 1H, Movie S1). Over ~2–10 min (Fig 1H), the force reached and exceeded the trap maximum of ~65 pN and pulled the bead out of the laser trap (Movie S1). This showed that in the presence of ATP, the ESCRT module can exert forces that reshape membranes.

We sought to determine which components of the ESCRT module were required for force generation (Fig. 2A). When Vps2 or Vps24 were the only ESCRT-III subunits, essentially no force was generated (Fig. 2D, E). Omission of Vps2 or Vps24 led to little or no force generation (Fig. 2F, H), consistent with the role of Vps2 in coupling of ATP hydrolysis to ESCRT-III remodeling, and a role of Vps24 in co-polymerizing with Vps2. When both Vps2 and Vps24 are present, a force rise up to 12 pN was produced, consistent with the ability of Vps24 and Vps2 to co-polymerize (*10*, *20*) (Fig. 2G). The inactivated mutant E233Q of Vps4 (*17*) failed to generate force (Fig. 2J). Deletion of the Vps4-binding MIM1 motif of Vps2 (Fig. 2D) (*18*, *19*) also abolished force generation. Thus, the ability of the ESCRT module to exert forces on nanotubes correlates closely with the presence of all the components that are crucial for ESCRT-mediated membrane scission and their individual integrity.

**Fig. 2.**
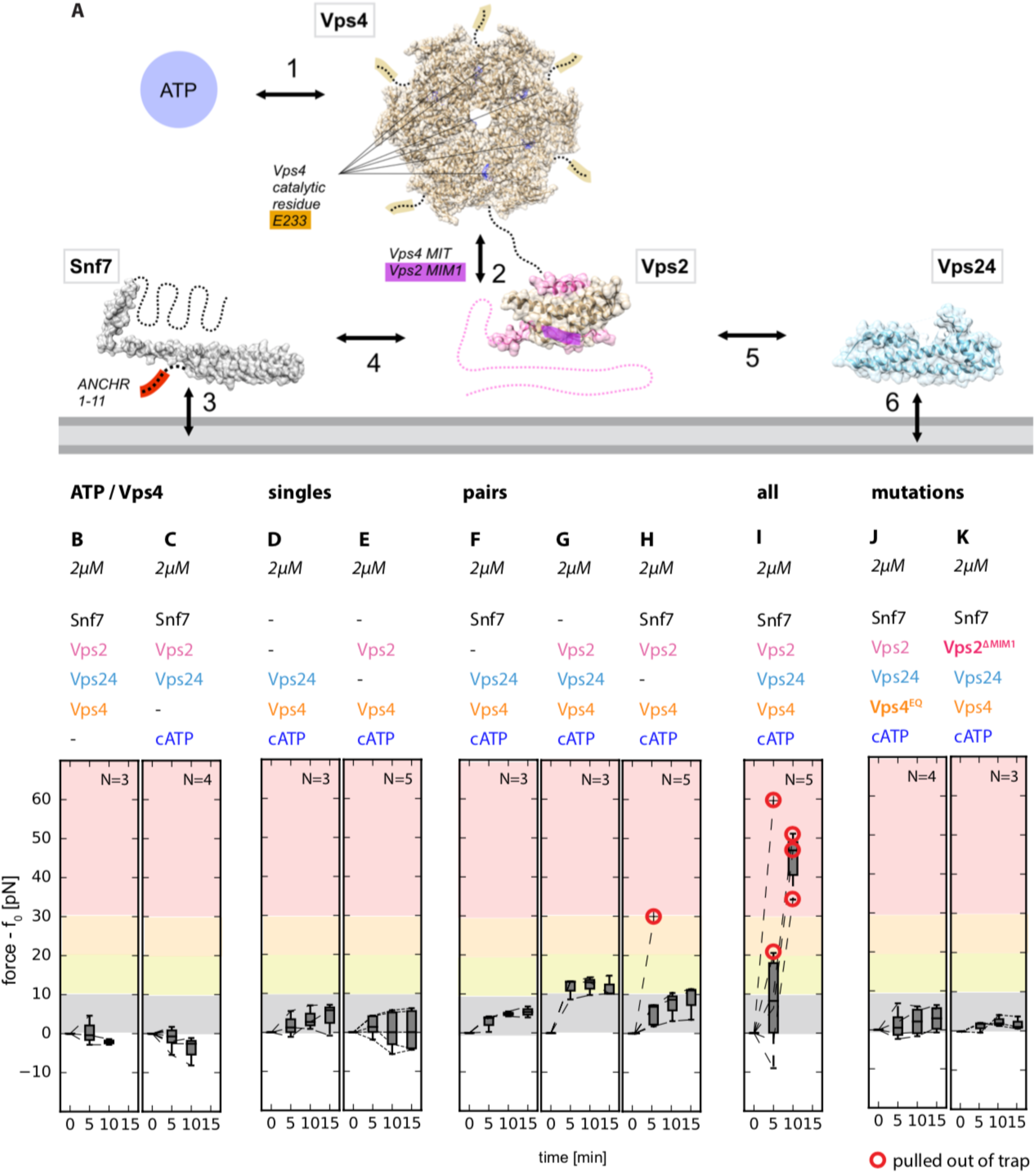
Molecular determinants of force generation. A. Interaction network of the ESCRT module. ESCRT proteins (filled-space structures and dashed lines) interact with the membrane (grey, bottom) as well as with each other. Key components (ATP), catalytic sites, and interacting motifs are highlighted in colors. B-E. Individual components of the module contribute differently to the force exerted on membrane tubes: ATP/Vps4 are essential for force generation (B, C), so is ATP hydrolysis (J, catalytically dead Vps4EQ mutant). Only the full module (I), or pairings of Vps2+Vps24 and Snf7+Vps2 (G, H), lead to significant force generation. Our data confirms known interactions within the ESCRT module and further points to additional interactions (arrows 4–6 in A) underpinning the activity of ESCRT machinery (see text).

Even 1200 sec after UV illumination, nanotubes containing ESCRT-III only (i.e. without Vps4 and ATP), or nanotubes containing the complete ESCRT module at the super-physiological concentration of 2 µM, remained intact and resistant to scission (Fig. 3A, B). Overexpression of ESCRT-III proteins in cells is known to generate dominant negative effects, so we lowered the concentrations of each component to 200 nM, similar to those estimated in yeast cells (*6*). 50% of the membrane nanotubes (n = 10) were severed within 1200 sec of ATP photo-uncaging (Fig. 3C). Forces ranging from ~6 – 40 pN were associated with the scission events (Fig. 3D). These data show that the ESCRT module reconstituted at physiological concentrations is capable of severing membrane nanotubes in the presence of ATP, and that scission is preceded by ESCRT-mediated force generation. Snf7 with its membrane-inserting ANCHR motif deleted (*29*) was ineffective, consistent with its loss of biological function (Fig. 3E).

**Fig. 3.**
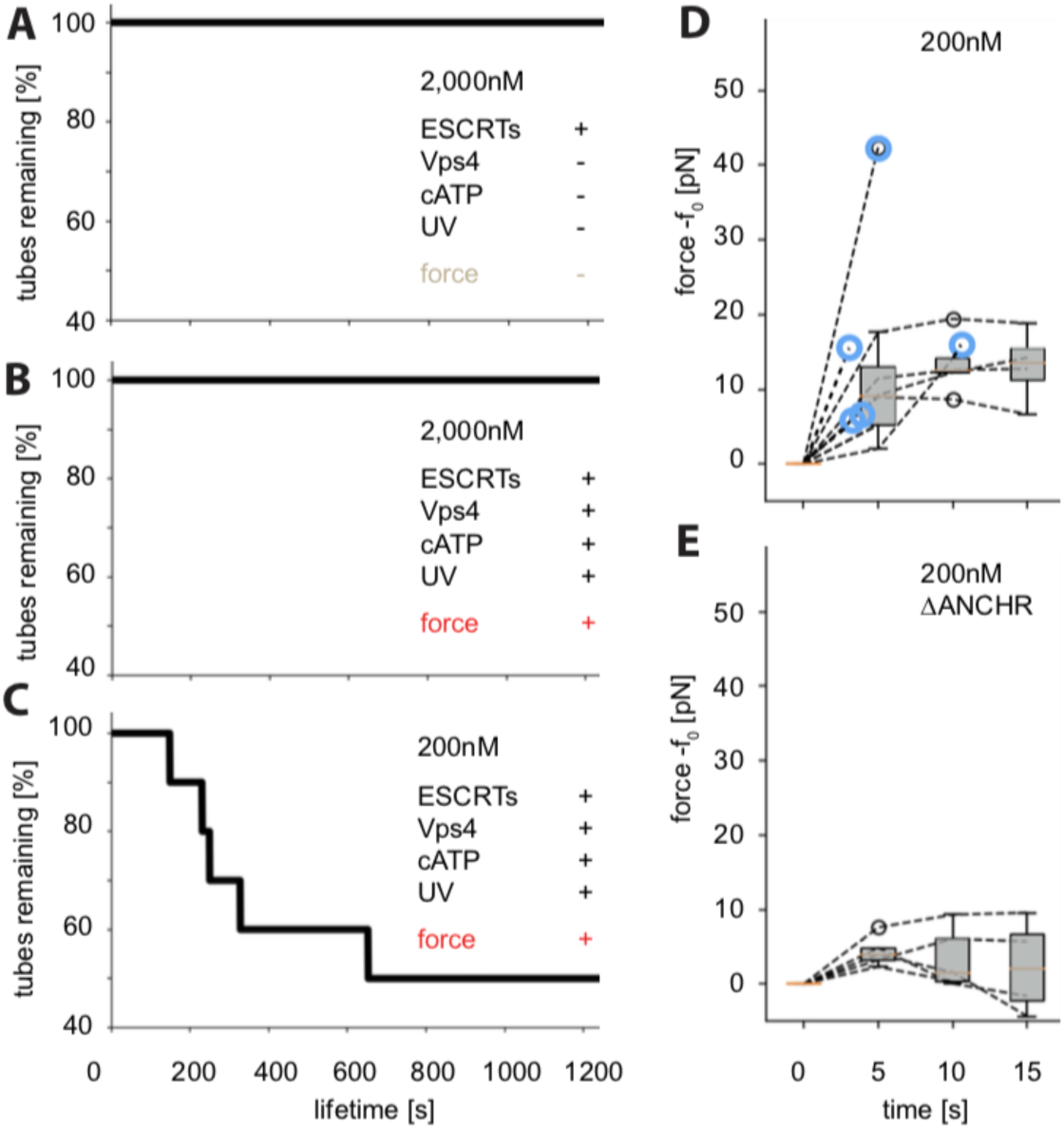
ATP-dependent membrane tube scission by ESCRTs at near physiological concentration. A. Membrane nanotubes remain stable (lasting at least 20 min; N=10) in control experiments where either ATP or Vps4 is knocked out. B. At high protein concentrations (2µM) ATP uncaging leads to force exertion (Fig. 2I), but the membrane tubes last with no scission occurring. C. When the full ESCRT module is encapsulated at near physiological protein concentrations (200nM), ATP uncaging consistently leads to both force exertion and tube scission on short timescales. D. Temporal profile of the force rise recorded under scission-promoting experimental condition. Blue circles indicate time points of a scission event. E. Temporal profile of the force in control experiments where the Snf7 ANCHR motif has been removed, and no scission is observed.

We integrated a confocal microscope with optical tweezing capability to image membrane nanotubes pulled from GUVs containing fluorophore-labeled ESCRTs (Fig. 4A–F, Movie S2). As compared to the activity with unlabeled proteins, scission occurred at somewhat shorter times, 100–200 s vs 200–300 s (Fig. 4G). Given that ESCRT-III polymerization is highly sensitive to the rate of nucleation (*30*), it is possible that the faster ESCRT-mediated scission represents multiple nucleation events. To maximize the signal in these experiments, Snf7 was labeled with the photo-stable dye, Lumidyne-550, and imaged with a resonant scanner and a GaAsP detector. We quantitated Snf7, Vps2, and membrane intensity using Gaussian fitting to the tube profile (Fig. 4H–M). The intensity in the membrane channel was nearly constant, with a drop of no more than 10% in the tube (Fig. 4L) and there was no change in the GUV size nor membrane intensity (Fig. 4M) preceding scission. These observations suggest that the force generation was not associated with a large-scale change in tube radius, and rules out that tube severing is the result of a change in the surface area of the GUV.

**Fig. 4.**
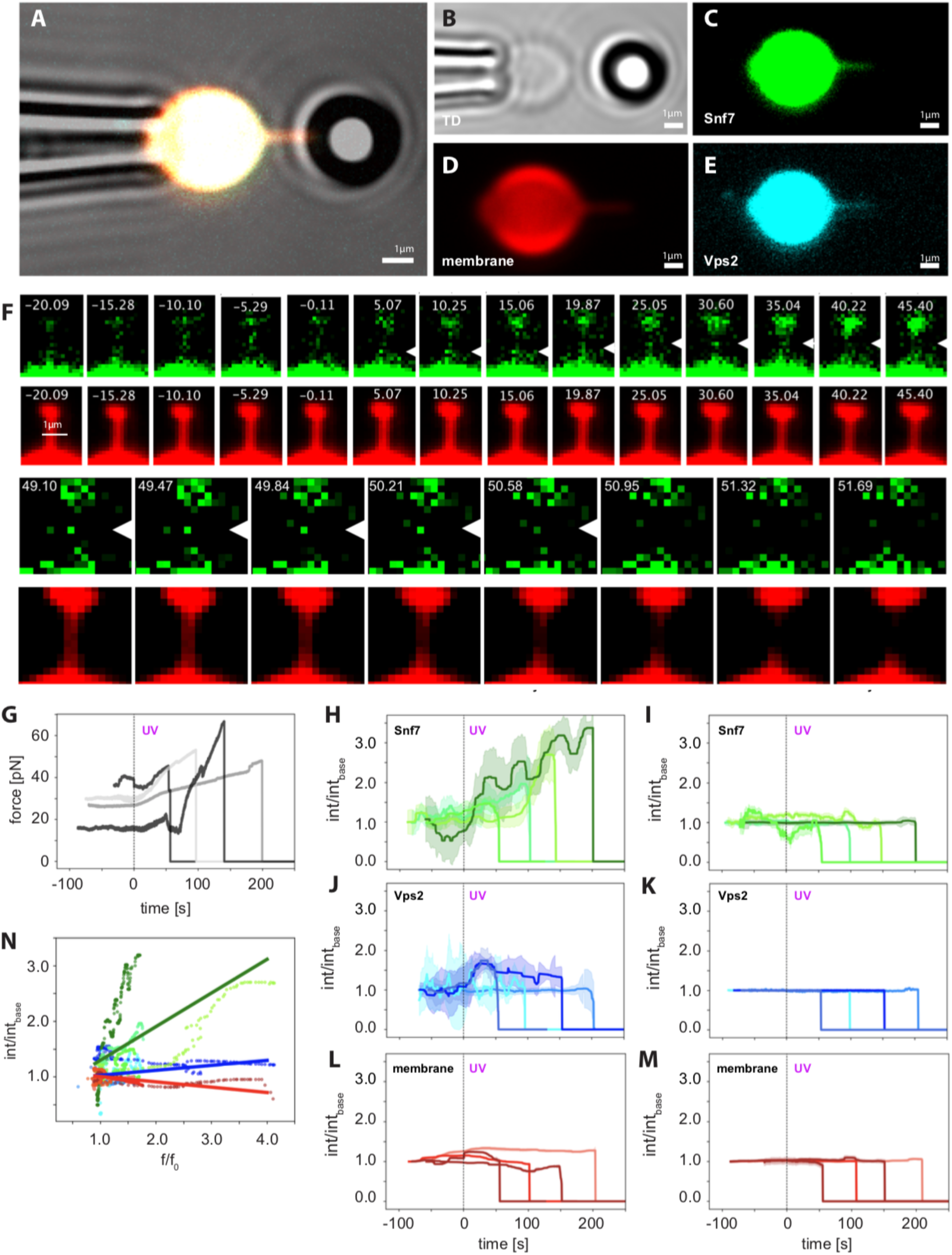
Confocal imaging and ESCRT-mediated membrane scission. A. Micrograph of the experiment setup: the GUV (center, white) encapsulating the ESCRT module and caged ATP is aspirated by a micropipette (left). A membrane nanotube has been pulled from the GUV using a bead (right) in an optical trap. Membrane, Snf7, and Vps2 are respectively labeled with different fluorophores (individual channels are shown in B-E). The merged channel (A) appears white. Scale bars: 1µm. F. Progression of a membrane scission event. UV illumination at t = 0 s. Forming Snf7 puncta are highlighted (white arrows) G, Force profile of 3 individual scission events. The characteristic tube pulling force signature is first observed at the beginning of each experiment (i.e. t = −150 to –70 s, left portion of the plot). A baseline force is then recorded (t = −70 to 0 s) before UV illumination. ATP released by UV uncaging leads to a rise in force and eventually scission (indicated by the drop of force to 0 pN). H-M, Normalized (and bleaching-corrected) fluorescence intensity profiles of the scission events for the tube (H, J, L) and the vesicle (I, J, K) in the three fluorescence channels (Snf7: H, I; Vps2: J, K; membrane: L, M; ± stdev shown as shaded area). N. Snf7 showed the largest correlation between the normalized tube fluorescence intensity (y-axis) and the normalized force (x-axis) observed during scission. Experimental data (circles) and a linear fit (lines) are color-coded to match the respective channels (Snf7: green, Vps2: blue, membrane: red).

The total fluorescence intensity from Snf7 in the tube increases prior to scission (Fig. 4J), with a close correlation seen between Snf7 intensity and the rise in force generation (Fig. 4N). Snf7 intensity in the GUV, however, is essentially unchanged (Fig. 4). Vps2 intensity also increased in the tube but to a lesser degree than Snf7 (Fig. 4J) whilst remaining constant in the GUV (Fig. 4K). The constant Snf7 intensity on GUV observed in our reverse-topology setup contrasts with a previous observation of a time-dependent increase of Snf7 recruitment to the exterior of highly acidic GUVs (*9*), which led to membrane stiffening and to force generation by an indirect mechanism. We can rule out such a bulk stiffening mechanism in our system given the lack of recruitment of Snf7 to the GUV membrane and the lack of correlation between GUV Snf7 intensity and force generation. On the other hand, the intensity of Snf7 in the tube correlates closely with the force rise (Fig. 4N). These data show that scission correlates with increased force and increased Snf7 in the tube.

Individual diffraction-limited puncta of Snf7 intensity appeared and disappeared within the tubes, as illustrated in Fig. 4. For example, two puncta shown in the upper row of Fig. 4F had a lifetime of ~10 and 15 s, respectively. A third punctum appeared at 49 s and its position corresponded to the site of membrane scission at 50 s. While not all of the Snf7 puncta formed in the tubes lead to scission, scission itself correlates with the appearance of a Snf7 punctum. We estimated an upper limit of ~170 Snf7 molecules were present in the nanotubes at the time of scission (Fig. S4). The number of Snf7 molecules at the scission site itself thus seems likely to be no more than a few tens of molecules.

It has generally been inferred that the core ESCRT-III proteins Snf7, Vps24, and Vps2, together with Vps4, comprise the minimal ATP-dependent scission machinery (*1*, *4*, *31*, *32*). Here, we directly confirmed this idea by visualizing scission in a minimal system that replicates a wide range of biologically validated structure-function relationships.

The most striking finding from the reconstituted system is that the core ESCRT-III proteins and Vps4 together exert an ATP-dependent axial force on the nanotube before severing. It had previously been proposed (*8*) and then demonstrated (*9*) that Snf7 filaments have a preferred curvature and may exert forces when bent above or below their preferred value. It had also been hypothesized that breakage or remodeling of ESCRT filaments by Vps4 could contribute to force generation (*4*, *15*, *20*, *33*). Our observations now provide the first experimental confirmation that ESCRTs indeed can generate force from within a narrow membrane tube, and show that this force is correlated with reverse-topology membrane scission.

ESCRTs (*7*, *10*, *11*) and the “normal topology” scission factor dynamin (*34*) have been visualized as cylindrical membrane coats, and Snf7 has also been seen in the form of large spirals of hundreds to thousands of copies (*9*). In the case of dynamin, extended coating of the tube is not needed, and one or a few rings appear capable of mediating scission (*35*). Our measurements of scission by diffraction-limited puncta of Snf7 are consistent with imaging in cells (*36*) and with scission by a ring of molecular dimensions, not an extended coat nor a micron-scale spiral. In fact, high concentrations of ESCRTs actually inhibit scission, suggesting that coat formation could antagonize scission. We also found that scission by ESCRT-III and Vps4 can occur mid-tube, contrary to past speculation that scission might be favored at saddle points or tube-vesicle junctions (*4*). Localization to specific loci within or at the ends of tubes may be governed by upstream factors such as ESCRT-I and ALIX.

Some models for membrane scission postulate that insertion of hydrophobic wedges into the proximal leaflet of the membrane disrupts lipid organization and promotes scission (*37*). In the context of the ESCRT system, this would be consistent with our observation that the hydrophobic ANCHR motif of Snf7 is important for scission (*29*). The mechanical nature of constriction and force generation remains to be elucidated through structural approaches. The ESCRT experimental system we developed here, both the reconstituted biochemical assay and the specifically tailored optical instrumentation, will make it possible to determine whether and how these upstream proteins might also participate in modulating the physical process of membrane scission.

## Acknowledgments

We thank J.-Y. Lee, H. Aaron, S. Ruzin and D. Schichnes for assistance with imaging, M. Vahey, D. Fletcher, P. Lishko and A. Roux for advice on the aspiration pipette set-up, A. Lee for assistance with the optical trap force calibration and C. Glick for advice with the microfluidics.

## Funding

Research was supported by a Marie Skłodowska-Curie postdoctoral fellowship ‘smStruct’ to J. S., an NSF predoctoral fellowship to M. R. P., and NIH grant R01AI112442 to J.H.H.

## Author Contributions

Conceptualization, J.S., M.R., D.H.G, S.Y., A.H.B., C.B., G.H. and J.H.H; Methodology, J.S., S.Y., M.R., D.H.G, A.H.B. I-H. L. and L-A.C.; Software, J.S.; Formal Analysis, J.S., A.H.B.; Investigation, J.S., S.Y., M.R., A.H.B. and D.H.G; Resources, J.S., S.Y., M.R., M.R.P., I-H. L., L.A.C. and X.R.; Data Curation, J.S; Writing – Original Draft, J.S. and J.H.H; Writing – Review & Editing, J.S., S.Y., M.R., M.R.P., I-H.L, L-A.C, A.H.B., D.H.G., X.R., G.H., C.B., J.H.H; Visualization, J.S.; Supervision, G. H, C.B. and J.H.H

## Competing interests

Authors declare no competing interests.

## Data and materials availability

All data is available in the main text or the supplementary materials. Code is available from the authors upon request.

## Supplementary Materials

Materials and Methods Figures S1-S4 Movies M1-M2 References (37–39)

